# Neuronal-immune Axis Alters Pain and Sensory Afferent Damage During Dental Pulp Injury

**DOI:** 10.1101/2022.12.23.521695

**Authors:** Ozge Erdogan, Benoit Michot, Jinya Xia, Lama Alabdulaaly, Pilar Yesares Rubi, Isaac Chiu, Jennifer L. Gibbs

## Abstract

Dental pulp tissue is densely innervated by afferent fibers of the trigeminal ganglion. When bacteria cause dental decay near the pulpal tissue, a strong neuronal and immune response occur, creating pulpitis, which is associated with severe pain and pulp tissue damage. Neuro-immune interactions have the potential to modulate both the pain and pathological outcome of pulpitis. We first investigated the role of the neuropeptide calcitonin-gene related peptide (CGRP), released from peptidergic sensory afferents, in dental pain and immune responses by using calca knock out (calca^−/−^) and wild type (calca^+/+^) mice, in a model of pulpitis by creating a mechanical exposure of the dental pulp horn. While CGRP did not contribute to facial mechanical hypersensitivity, at an early time point, it did contribute to spontaneous pain-like behavior. We also found that CGRP contributed to recruitment of neutrophils and monocytes, while not clearly affecting the progression of pulpal pathology histologically. When we depleted neutrophils and monocytes, we found that there was more sensory afferent loss, tissue damage and deeper spread of bacteria into the pulp tissue, while there was a reduction in facial mechanical hypersensitivity compared to control animals at a later time point. Overall, we showed that there is a crosstalk between peptidergic neurons and neutrophils in the pulp, modulating the pain and inflammatory outcomes of the disease.

## INTRODUCTION

The pulp tissue resides within the hard tissues of a tooth. Upon injury, such as bacteria forming deep caries, pulp tissue initiates a robust, often very painful, inflammatory response, the state which is referred to as pulpitis [28]. Clinically, the degree of tissue damage during pulpitis, including the die-back of sensory afferents, is often determined to be irreversible based on the presence of painful symptoms, leading to invasive treatment choices including root canal treatment or tooth extraction [28]. However, the relationship between the severity of pain and degree of tissue damage, including sensory afferent loss, during pulpitis is not clear [11]. Studying the mechanisms of neuro-immune interactions and tissue damage, we can have a better understanding of the relationship between pain and pathological outcomes of pulpitis.

Dental pulp tissue is highly innervated [9; 20]. On average, each molar tooth in the mouse is innervated by fifty neurons [9]. The types of neurons innervating dental pulp are diverse and include peptidergic afferents expressing the neuropeptide calcitonin gene-related peptide (CGRP), which compromise about 30% of all pulp neurons [9; 15; 39]. CGRP mediates headache and trigeminal pain both through peripheral and central mechanisms within the trigeminal system [8; 23; 29; 54; 57]. CGRP expression is also higher in the pulp of patients with severe dental pain compared to non-painful cases [1]. However, whether interfering with CGRP signaling from sensory afferents affects pain outcomes in pulpitis is not well established [3; 12]. In addition, CGRP released from peptidergic neurons modulates innate immune cell activity through its cognate receptor receptor associated modifying protein (RAMP1) and Calcitonin receptor-like receptor (Calcrl) expressed by these innate immune cell types in different barrier tissues [6]. For example, CGRP has been shown to modulate neutrophil recruitment and antimicrobial signaling in a mouse model of *S. pyogenes* skin and soft tissue infection [41]. Ablation of nociceptors and blockade of CGRP signaling led to increases in neutrophil recruitment, which resulted in decreased dermonecrosis [41]. However, whether sensory neuron derived CGRP regulates innate immune cells in dental pulp is unknown.

Neutrophils and monocytes are crucial players of innate immune response, serving as the first line of defense against tissue injury and pathogen invasion, being equipped with different antimicrobial mechanisms [49; 50]. Neutrophils are also critical for clearance of tissue debris and wound healing [25]. Pain is coupled to both the innate immune response and wound healing [38; 42; 44; 51]. Innate immune cells release multiple proinflammatory cytokines such as IL-1β and TNF-α, which can sensitize nociceptors to drive inflammatory and neuropathic pain [18; 35; 42; 55]. On the other hand, neutrophils can be important for suppression of long-term development of chronic pain: in the Complete Freund’s adjuvant (CFA) inflammatory pain model, it has been shown that while withdrawal threshold was initially higher in animals where neutrophils were depleted, for these animals it took them significantly longer to return to baseline withdrawal thresholds [38]. Investigating the role of neutrophils/monocytes in dental pain and sensory afferent damage can improve our understanding of relationship between pain, innate immunity, and pathological outcomes during pulpitis.

## MATERIALS AND METHODS

### Animals

All animal experiments were approved by the Institutional Animal Care and Use Committee (IACUC) at Harvard Medical School. C57BL/6 mice were purchased from Jackson (Jax) Laboratories (Bar Harbor, ME). *Calca-GFP-DTR* [31] and *Calca*^*-/-*^ *and Calca*^*+/+*^ [37] mice were bred in house (Harvard Medical School). Age-matched 7-to 12-week-old littermate mice of both genders were used for experiments. For pain behavior assays, only female mice were used.

### Dental Pulp Injury Procedure

Clinically, as dental caries progresses by degradation of dental hard tissues, the pulp tissue gets exposed to caries microbiome and oral environment and the process of progressive tissue degradation begins, the stage referred as pulpitis. Dental pulp injury, drilling of hard tissues of the tooth to expose the pulp to oral environment and bacteria is a model that has been widely used to model pulpitis [16]. To perform, dental pulp injury, mice were anesthetized via intraperitoneal injection of 100 mg/kg ketamine + 10 mg/kg xylazine in sterile phosphate buffered saline (PBS). The pulp exposures were performed in accordance with published protocols [16; 27]. Briefly, the maxillary first molar was drilled with a ¼ round bur at low speed in a way that only the middle pulp horn was drilled until the pulp was exposed which allowed for bacteria inoculation and penetration through the injured sited (Supplementary Figure 1). The contralateral maxillary first molar served as the uninjured control. Sham mice received the same anesthesia and had their mouths held open in a similar manner and duration. For pain behavior and histology experiments, only the maxillary first molar for each animal was injured. For flow cytometry experiments, maxillary first, second and mandibular first molars were injured on each animal and teeth from 5 mice were pooled. All procedures were performed by one investigator (OE).

### Gr-1 Neutrophil/Monocyte Depletion

Mice received 250μg Gr-1 antibody (anti-mouse Ly6G/Ly6C (Gr-1), clone RB6-8C5, BioCell) two hours before the dental pulp injury procedure and on day three, while the control mice received 250μg isotype control antibody (rat IgG2b isotype control, anti-keyhole limpet hemocyanin, BioCell).

### Spontaneous Pain-like Behavior Measured by the Mouse Grimace Scale

The mouse grimace scale is validated for capturing spontaneous pain-like behavior in mice [21; 26; 45; 58]. Animals were acclimated in clear acrylic chambers (8 W x 8 H x 8 L cm) a day prior to baseline testing for 20 minutes. Animals were videotaped for 10 minutes at baseline and selected time points (day1/2, day3/4, day6/7) with individual cameras (GoPro, Inc). The first image with a clear view of the animal’s face of every minute of the video was extracted using iMovie (Apple, Inc), in total leading to 10 still image per video. Scoring of the images was performed by investigators blinded to treatment allocation and timing relative to injury. As previously described for each image, orbital tightening, nose bulge, cheek bulge, ear position were scored (0 “not present”, 1 “moderately visible” and 2 “severely visible”). Because whiskers were not clearly visible in some images, whisker scoring was excluded. A mouse grimace scale score for each image was then obtained by the averaging of given scores to each facial features. The final score for each video was calculated by averaging of scores of 10 images.

### Mechanical Sensory Response Measured by Facial Von Frey Stimulation

For this assessment, animals were placed in modified confined chambers with adjustable openings as previously described [45]. The animals would put their face out of the opening but because they were elevated from the floor, they also did not intend to escape. In this way, we were able to perform the facial stimulation using Von Frey filaments. The maxillary first molar teeth are adjacent to the periorbital area beneath the eyes, mechanical stimulation was performed there, ipsilateral to the injury by blinded single investigator (O.E.). The 0.008, 0.02g, 0.004, 0.07g, 0.16g, 0.4g filaments were tested at baseline and at selected time points (day 1, day 3, day 7) once for experiments performed with *Calca*^*-/-*^ *and Calca*^*+/+*^ mice. For Gr-1 depletion experiment, based on our data from *Calca*^*-/-*^ *and Calca*^*+/+*^ experiments, we only used 0.02g (non-painful stimulus), 0.07g (allodynia response stimulus) and 0.16g (painful stimulus) filaments once as the mechanical response score differences between injured and non-injured animals were clear with these filaments. The animals’ response to the stimulus on the face was scored by using a scoring system slightly modified from the previous publication in which the same experimental set up was used [45]. We calculated an overall mean mechanosensory response score by averaging the scores given by the mice to each filament for each time point. All measurements were performed by one investigator (O.E.).

| Score | Response |
| --- | --- |
| 0 | no response |
| 1 | orientation to the stimulus or a slower head turn away from the stimulus |
| 2 | a rapid withdrawal |
| 3 | rapid withdrawal followed by facial wipes or when mice stop taking the head out |

### Pulp Tissue Cell Suspension Preparation

Animals were humanely euthanized by CO_2_ inhalation. Maxillae/and mandibles were dissected, put in culture media on ice. Any tissues surrounding the teeth were removed. The crowns of teeth were separated at the crown-root interface which allowed both for exposure and collection of coronal pulp tissue. The molar crowns with pulp tissues were then incubated in digestive solution (4mg/ml collagenase and 4mg/ml dispase II) at 37 °C for 1 hour. Later, the tissues within the digestive solution were pipetted vigorously for the pulp tissue to be detached from dentin. The samples were spun down at 300g for 5 minutes, washed with flow buffer (2% FBS, 1μM EDTA, 0.1% sodium azide) and filtered using 70 μm mesh.

### Splenocyte Cell Suspension Preparation

Spleens were collected in culture media on ice and were homogenized using 70 μm cell strainer and by using culture medium. The cell suspension was washed using and splenocytes were centrifuged at 1500rpm, at 4 °C for 5 minutes and resuspended in red blood cell lysis buffer for 10 minutes. The splenocyte cell suspension was washed once again, and finally cells were resuspended in culture media.

### Flow Cytometry

The cell suspension was spun down at 300g for 5 minutes and resuspended in flow buffer. The cell suspension was first incubated with FcR blocking reagent (Miltenyi Biotec, 130-092-575) for 10 minutes, washed with flow buffer and then incubated with the following antibodies (Zombi Aqua-KO525 (1:1000, Biolegend), CD45-Cy7 (1:100, Biolegend), CD11b-PE (1:100, Biolegend), Ly6G-APC (1:100, Biolegend), Ly6C-AF700 (1:100, Biolegend), CD3-FITC (1:100, Biolegend), CD4-PE-cy7 (1:100, Biolegend), CD8-perpCy5.5 (1:100, Biolegend), F4/80-PB (1:100, Biolegend) for 30 minutes. Cell suspensions were washed again, centrifuged, resuspended in 1% paraformaldehyde (PFA) and filtered through a 40 mm mesh. Counting beads (Biolegend) were added to each sample. Flow cytometry was performed on a CytoFLEX flow cytometer (Beckman Coulter Life Sciences) and data was analyzed using FlowJo software (FlowJo LLC).

### Tissue Processing for Histology

Animals were perfused with PBS with heparin, followed by 4% PFA. Maxillae were dissected. All tissues were post-fixed with 4% PFA overnight. Decalcification of maxillae using 18% EDTA at 4°C was performed for 3-4 weeks. For paraffin embedding, tissue processing including dehydration process was performed using a tissue processor (TP1020, Leica). Maxillary jaws were then embedded with paraffin and sectioned (6μm). For all the other immunofluorescence staining experiments, the maxillary jaws were dehydrated by 30% sucrose for one day, followed by 50% sucrose for one day at 4°C, then embedded in optimum cutting temperature compound (OCT) for frozen tissue storage and sectioning, at 30μm thickness.

### H&E Staining

Rehydration, hematoxylin & eosin staining, dehydration and mounting of the slides were completed. Sections of the coronal pulp were examined by a board-certified oral and maxillofacial pathologist, blinded to treatment allocation (L.A.). Injured (middle), non-injured (mesial and distal) coronal pulp horns were investigated and scored individually for extent necrosis (0-3) and extent of odontoblast loss (0-3) (0: no changes, 1: 1/3 of the area of interest is affected, 2: 2/3 of the area of interest is affected, 3: 3/3 of the area of interest is affected) and edema (0: absent, 1: present). The overall score for each section was calculated by averaging mesial, distal, and middle pulp horns. Three sections per animal were scored and averaged, to obtain the individual animal score.

### Immunofluorescence Staining

Cryosections were washed with PBS. Sections were blocked with 5% goat serum and 1% triton for 1 hour at room temperature. Sections then were incubated with the primary antibodies overnight at 4°C. Primary antibodies were: rabbit anti-beta tubulin (tuj1) (1:500, abcam), rat anti-Ly6g/Ly6C (1:200, abcam), rat anti-F4/80 (1:200, abcam). Sections were then washed with 1% goat serum and 1% triton and were incubated with the secondary antibodies for 2 hours. Secondary antibodies were: goat anti-rabbit Alexa fluor 546 (1:500, ThermoFisher), anti-rat Alexa fluor647 (1:500, ThermoFisher). Image acquisition was performed by confocal microscope (Leica, Stellaris). Equal number of stacks were imaged using the same acquisition settings. Image processing was performed using Fiji [48] with the same settings across experiments. To quantify the area with sensory afferent loss, we divided the pulp chamber in two areas as: 1-injured middle pulp horn with the connection parts to non-injured pulp horns 2-non-injured pulp horns. For both areas, we first measured the whole area stained with dapi, then the area without tuj1 staining and calculated the percent area with sensory afferent loss. Quantification of one section from each animal (5 from isotype, 5 from Gr-1 groups) was performed using Fiji.

### Fluorescence In-situ Hybridization

Sections were de-paraffinized using xylene and rehydrated using 100% ethanol. Half of the sections in one slide were incubated with 1:200 Cy3-congugated nonsense eubacterial probe (5’-/5Cy3/CGACGGAGGGCATCCTCA-3’) whereas the other half of the same slide was incubated with 1:200 Cy3-conjugated eubacterial probe (5’-/5Cy3/GCTGCCTCCCGTAGGAGT-3’) in 20% SDS, 1:10 formamide, and 9:10 bacterial hybridization buffer overnight in 50°C in tissue culture incubator. Sections were washed first with hybridization buffer followed by PBS before mounting.

### Statistical Analysis

Statistical comparisons of two groups for a single variable with normal distributions was analyzed by unpaired Student’s t-test. Statistical comparisons of three or more groups at a single time point were analyzed by one-way ANOVA with Tukey post-tests. Outcomes of pain behavior assays were analyzed by two-way ANOVA with repeated measures. Between subject effects were used to determine if there had been any significant effects of time, experimental group, or a significant interaction, followed by with post-hoc Dunnett’s and Sidak tests where necessary. A p-value less than 0.05 was considered statistically significant. Statistical analyses were conducted using Prism 9 (GraphPad Software, LLC) and Excel (Microsoft).

## RESULTS

### CGRP+ neuronal afferents reside at the dentin-pulp junction and die back as the disease progresses

We first wanted to capture the spatial distribution of CGRP+ fibers within the pulpal tissues in relation to all nerves of the pulp, marked by beta tubulin 3 (tuj1), and how the neuronal architecture was affected after the pulp injury. CGRP-α is the main isoform of CGRP expressed by nociceptive afferents [46]. Calca-GFP reporter mice have been used to study CGRP+ peptidergic sensory innervation of the skin and other barrier sites [31]. Using these mice and anti-beta tubulin III (tuj1) antibody to stain for all nerve fibers, we observed that CGRP+ fibers extend along the dentin-pulp junction through all three pulp horns in healthy, intact pulp tissue (Figure 1-A). On day 0, immediately after mechanical injury of the dental pulp, we observed minimal afferent loss located at the middle, injured pulp horn (Figure 1-B). On day 1, we observed continuity of Tuj1 expressing afferents along with CGRP+ fibers between the three pulp horns and damage was localized to the injury site (Figure 1-C). At day 3 and 7, the extent of the afferent die-back increased and we observed loss of the continuity of tissues connecting the pulp horns as well as partial loss of afferents within the other non-injured pulp horns (Figure 1-D, E). Qualitatively, we observed that especially on day 7, the loss of CGRP+ fibers was prominent, even in relation to all Tuj1+ nerve fibers (Figure 4-E).

**Figure 1.**
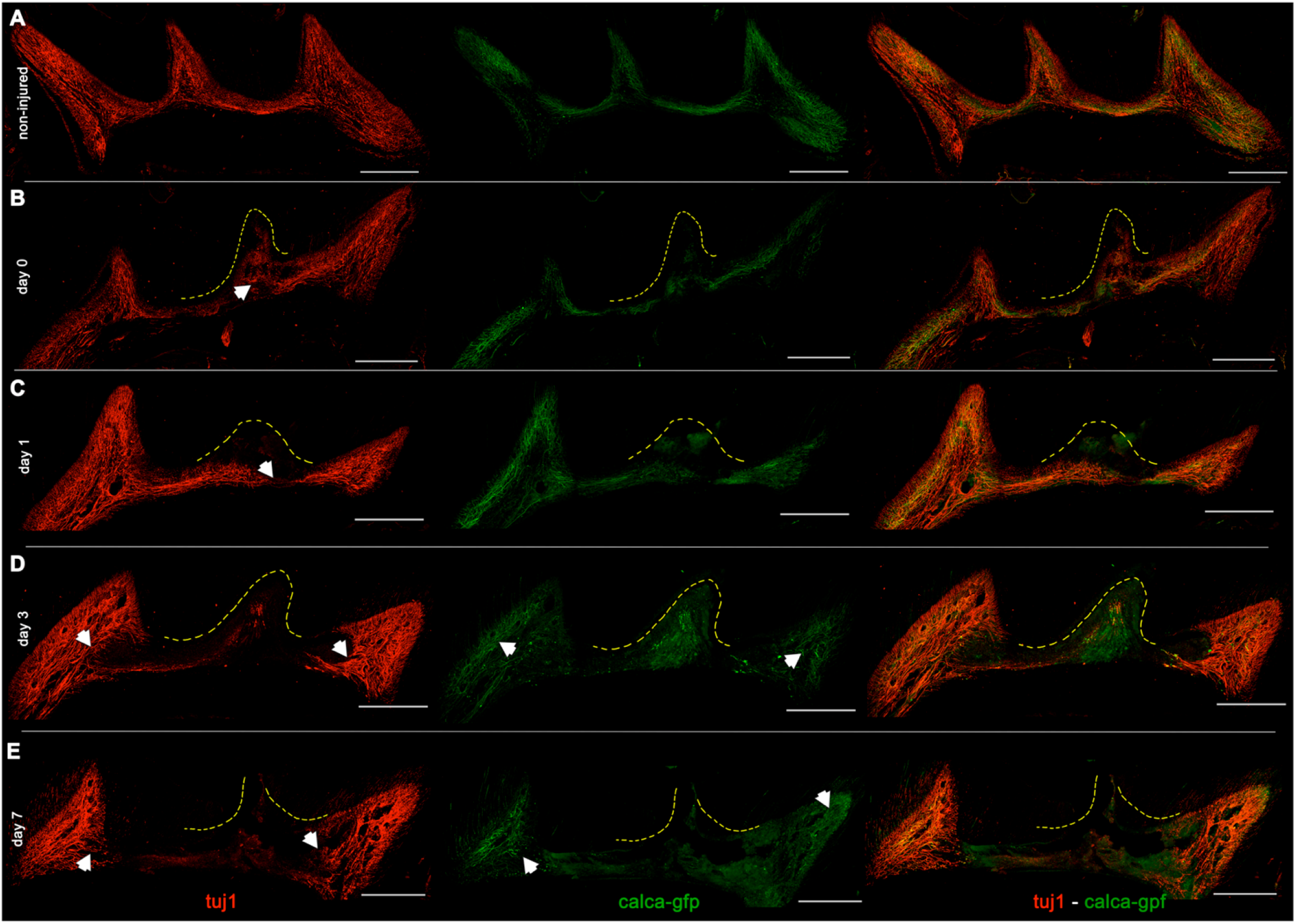
CGRP+ neuronal afferents are an important neuronal subpopulation of the pulp and die back as the disease progresses. A-Non-injured pulp. CGRP+ fibers extend along the dentin-pulp junction. B-Pulp, immediately post-injury. The arrow shows the presence of intact afferents within the injured middle pulp horn C-Pulp, one day after injury. The arrow points out the continuity of intact fibers just right after their interruption. Most of the CGRP+ fibers are also still present. D, E-Pulp, three and seven days after injury. The arrows point out the extent of die back by marking the border of remaining intact afferents within the non-injured pulp horns. CGRP+ fibers are scarcer, mostly present at the very distant ends of the pulp chamber. The middle pulp horn is the mechanically injured pulp horn in each image. The first column includes images showing beta tubulin III (tuj1) immunofluorescence, the second column shows images of CGRP+ fibers, traced by calca-gfp reporter mice, the third column shows co-localization of tuj1 with calca-gfp signal. Scale bars are 200 μm. Original magnification 20X.

### CGRP increases spontaneous pain-like behavior at day 1 but do not contribute to mechanical hypersensitivity

In order to assess the contribution of CGRP to pain during pulp injury, we investigated whether facial mechanical hypersensitivity responses and spontaneous pain-like behavior captured by grimace scoring was different in Calca^−/−^ mice compared to wild-type mice. We found that the mechanical response scores were similar for both genotypes across each time point in non-injured animals (Figure 2-A, B). Both Calca^−/−^ and Calca^+/+^ mice exhibited a higher mechanical response score at day 1, 3 and 7 after the injury compared to baseline measurements (p=0.015, p=0.015, p=0.001, p=0.01, p=0.090, p=0.001) (Figure 2-A, B), supporting that dental pulp injury produces mechanical hypersensitivity in the facial skin. However, we did not find any difference in mechanical response scores between Calca^−/−^ and Calca^+/+^ mice at any time points (Figure 2-A, C).

**Figure 2.**
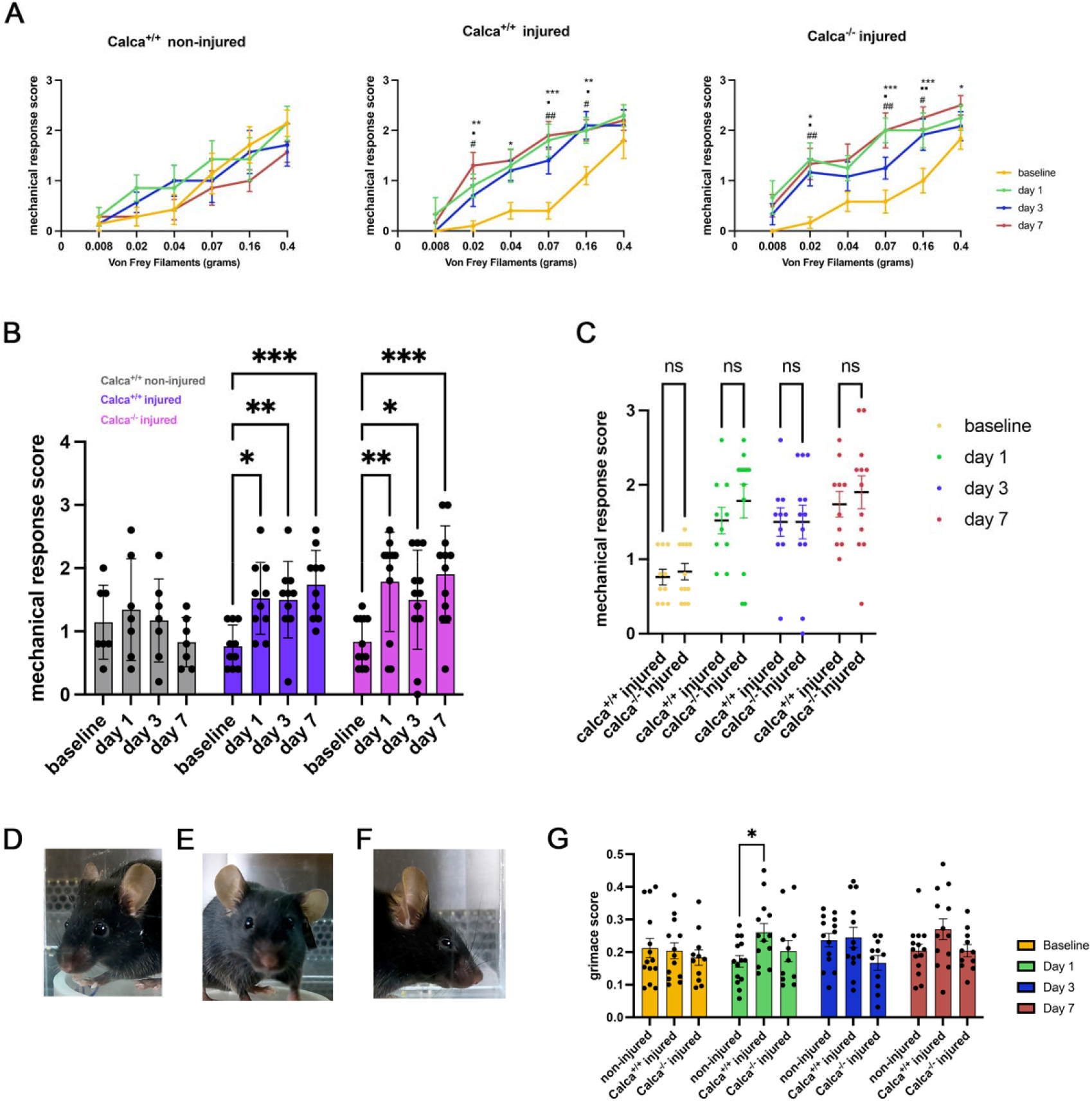
CGRP contributes to spontaneous pain-like behavior but not to mechanical hypersensitivity response after dental pulp injury. A-Plots present mechanical response score of non-injured (n=7; calca^+/+^ and calca^−/−^ grouped together), calca^+/+^ (n=10) and calca^−/−^ (n=12) animals to Von Frey filaments with different weights at different time points. B-In this plot, each data point represents the mean mechanical response score of one animal obtained by averaging the mechanical response scores given to each Von Frey filament. C-Comparison of mean mechanical response score of calca^+/+^ and calca^−/−^ animals. D, E, F-Representative extracted images of mouse used to examine facial expression changes for grimace scoring e.g. normal eyes, ears, cheek and nose (D), nose bulge, cheek bulge (E), eye squinting (F) G-Comparison of grimace scores between non-injured (calca^+/+^ and calca^−/−^ grouped together), injured calca^+/+^ and calca^−/−^ across different time points. A, B-Repeated measures of ANOVA with Dunnett’s multiple comparisons. A-# comparison of day 1 to baseline,▪ □ comparison of day 3 to baseline, * comparison of baseline to day 7. G-Two-way ANOVA with Dunnett’ s multiple comparison tests. #▪ □ ∗p < 0.05, ##▪ □▪ □ ∗∗p < 0.01, ###▪ □▪ □▪ □ ∗∗∗∗p < 0.0001. Mean ± SEM.

We also analyzed grimace scores as a measure of spontaneous pain-like behaviors in Calca^+/+^ and Calca^−/−^ injured animals and non-injured animals. There was statistically significant differences in grimace scores for Calca^+/+^ and Calca^−/−^ injured animals compared with non-injured animals (Two-way ANOVA: side: p=0.009, F=4.8, DOF=2; time: p=0.64 F=0.5, DOF=3; interaction: p=0.3, F=1.2, DOF=6) We found that, there was no difference at baseline among groups (p=0.96, p=0.64) (Figure 2-F). On day 1 after injury, grimace scores of wild type mice were significantly higher compared to non-injured animals (p=0.022) whereas there was no difference between non-injured animals and Calca^−/−^ mice (Figure 2-F). On day 7 after injury, we observed a similar trend, where grimace scores of wild type mice were higher compared to non-injured animals (p=0.11) whereas grimace scores of non-injured and Calca^−/−^ mice were similar (p=0.99) (Figure 2-F). These data suggest that CGRP may contribute to spontaneous pain-like behaviors due to dental pulp injury.

### CGRP contributes to recruitment of immune cells at day 1, but do not alter tissue damage

Next, we looked at whether CGRP modulated the innate immune response after pulp injury. One day after injury the pulp tissue was collected from Calca^−/−^ and Calca^+/+^ mice and flow cytometry analysis was performed. We found that CGRP contributed to recruitment of neutrophils and monocytes one day after injury. We found that there were fewer total CD45+ immune cells (p=0.011) in Calca^−/−^ mice compared to wild type mice after injury (p=0.017) (Figure 3-B, C, G). Among myeloid immune cells, we found that there were fewer CD11b+F4/80-Ly6G+neutrophils and CD11b+Ly6G-Ly6C+ monocytes in Calca^−/−^ mice compared to wild type mice (p=0.02) (Figure 3-D, E, G). While cell numbers differed, the overall proportions of immune cells in the dental pulp were similar between Calca^−/−^ and Calca^+/+^ mice (Figure 3-F). These data suggest that CGRP regulates overall recruitment of immune cells, of which the majority were neutrophils and monocytes. We then investigated whether the immune cell recruitment difference in Calca^−/−^ mice compared to wild type mice resulted in differences in the severity of tissue pathology. Since histological changes would not be detected at day 1, we investigated necrosis, odontoblast loss, and edema at day 7. There were no differences in tissue necrosis, odontoblast loss or edema between Calca^−/−^ mice compared to wild type mice at this time point (p=0.7, p=0.74, p=0.09, respectively) (Supplementary Figure 2).

**Figure 3.**
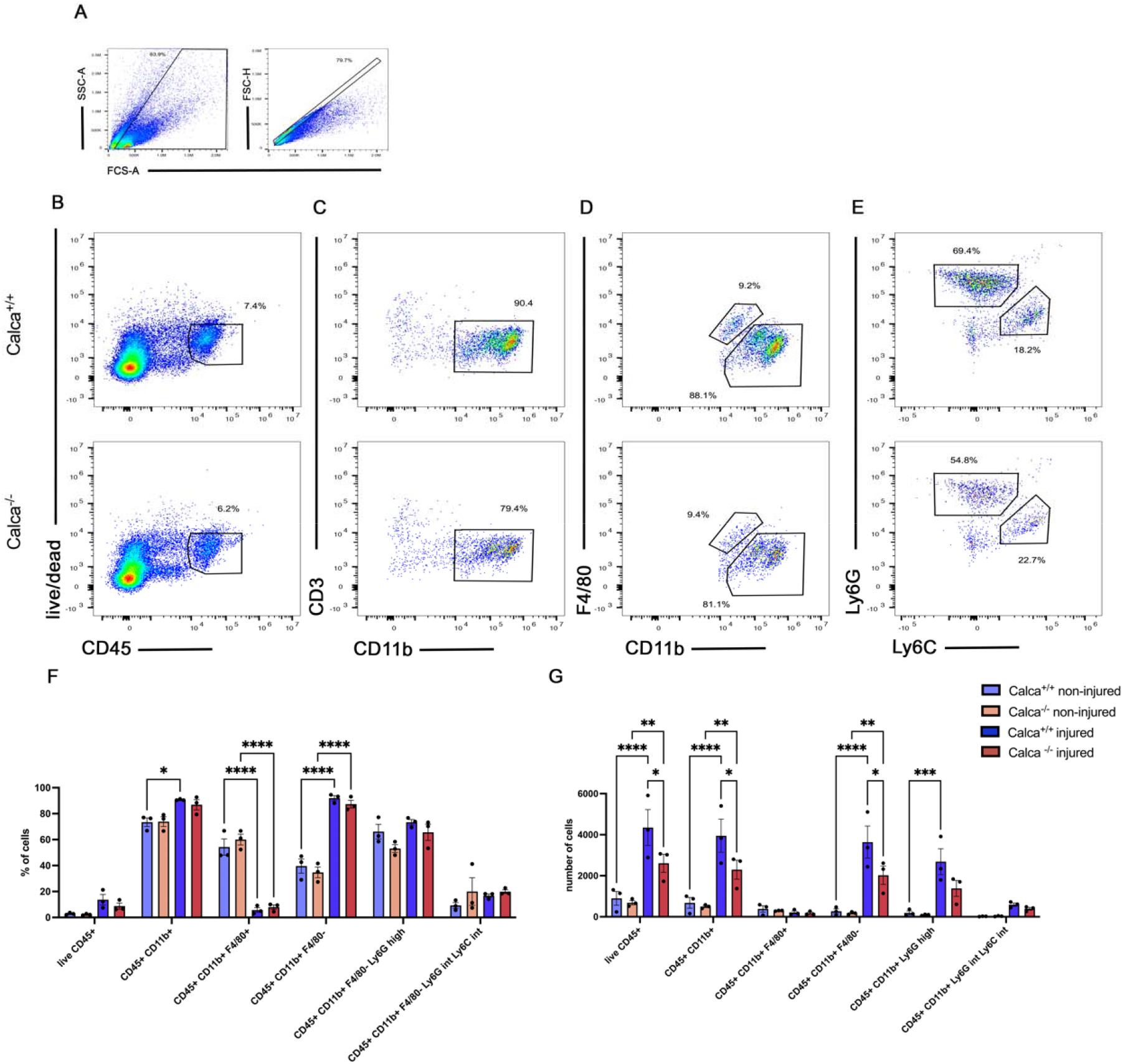
CGRP contributes to recruitment of neutrophils/monocytes at one day after injury. A-Representative flow cytometry (FC) plots showing initial steps of the gating strategy. B-Representative FC plots to gate CD45+ live immune cells D-Representative FC plots to gate CD11b+ myeloid cells E-Representative FC plots to gate F4/80+ macrophages F-Representative FC plots to gate F4/80-myeloid cells into Ly6Ghigh and Ly6Gint Ly6C int neutrophils/monocytes. The first row includes representative FC plots of calca^+/+^ animals and the lower row includes representative FC plots of calca^−/−^ animals. F-Percentage changes of immune cell populations plotted from three independent experiments of Calca^−/−^ and Calca^+/+^ animals. G-Number of immune cell populations plotted from three independent experiments of calca^−/−^ and calca^+/+^ animals. F, G-Two-way ANOVA with Sidak’s multiple comparisons test. ∗p < 0.05, ∗∗p < 0.01, ∗∗∗∗p < 0.0001. Mean ± SEM.

### Neutrophils are recruited into the dental pulp and invade denervated areas

Neutrophils are critically involved in bacterial clearance, tissue injury and wound healing. We next wanted to determine the kinetics of neutrophil recruitment during pulp injury, and how this relates with CGRP+ afferents. We utilized anti-Ly6C/6G to stain for neutrophils and monocytes. We found that in the non-injured healthy pulp, the presence of neutrophils/monocytes was scarce where intact CGRP+ fibers were visible (Figure 4-A). On day 1 after injury, we observed a very significant increase in neutrophils/monocytes at the injury site, where we concurrently observed the loss of CGRP+ fibers (Figure 4-C). On day 3, we observed that neutrophils and monocytes were extending towards the other non-injured pulp horns. We observed the presence of intact CGRP+ fibers were mostly outside of the area invaded by these immune cells (Figure 4-D). On day 7, the presence of neutrophils/monocytes seem to be more spread out within the pulp chamber and their rounded clear morphology became less distinct. At this time point, the presence of CGRP+ fibers were very scarce, mostly localized at the distant ends of the non-injured pulp horns (Figure 4-E).

**Figure 4.**
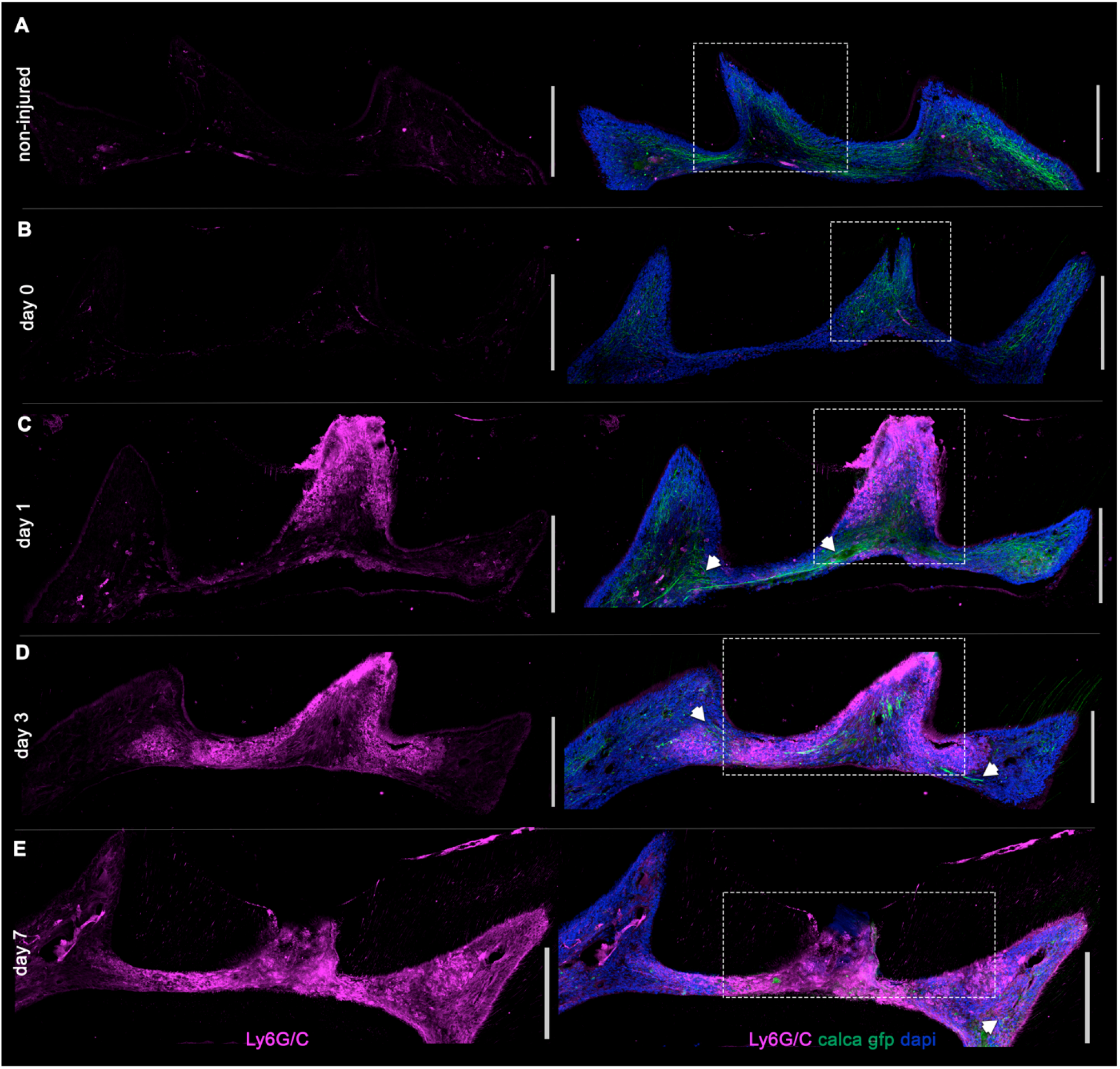
Spatial Investigation of neutrophils/monocytes with CGRP+ afferents within the pulp chamber. A-Non-injured pulp. The box shows the intact middle pulp horn with intact CGRP+ fibers and very few neutrophils/monocytes. B-Pulp, just right after injury. The box shows the injured middle pulp horn where most of the CGRP+ fibers are still intact and with very few neutrophils/monocytes. C-Pulp, one day after injury. The box shows the injured pulp horn densely populated by neutrophils and monocytes. The arrows point out the presence of CGRP+ fibers where there are fewer neutrophils/monocytes D, E-Pulp, three and seven days after injury. The arrows point out areas where the distinct CGRP+ fiber morphology is visible, just adjacent to but outside of where neutrophils and monocytes are densely present. The middle pulp horn is the mechanically injured pulp horn in each image. The first column includes images showing Ly6G/C immunofluorescence, the second column shows Ly6G/C co-localization with calca-gfp and dapi. Scale bars are 200 μm. Original Magnification 20X.

### Depletion of neutrophils and monocytes leads to more sensory afferent loss and further bacterial invasion

Because we observed that during pulpitis, the intact CGRP+ fibers were mostly located in areas that were not densely populated by neutrophils/monocytes, we aimed to investigate if continuous neutrophil and monocyte accumulation after initial recruitment contributed to tissue/sensory afferent damage in this enclosed environment of the pulp. We utilized Gr-1 antibody to deplete neutrophils/monocytes [4; 14]. We first confirmed the substantial reduction in the percentage of neutrophils and monocytes by flow cytometry experiment using splenocytes collected at day 6 (Figure 5-B). We found that at day 6 after injury, histologically, in the Gr-1 group, there was significantly more sensory afferent loss at the injured site (p=0.002) and at the non-injured pulp horns (p=0.028) (Figure 5C, D). In addition, there was more tissue necrosis and odontoblast loss (p=0.020 and p=0.011, respectively) and more edema formation in the Gr-1 group compared to isotype antibody injected control group (Figure 6-A). By performing an 16S FISH experiment, we found that after neutrophil depletion there was greater bacterial accumulation, extending to the non-injured pulp horns, whereas in isotype group, the bacteria remained localized to the injured area (Figure 6-B, C, D, E).

**Figure 5.**
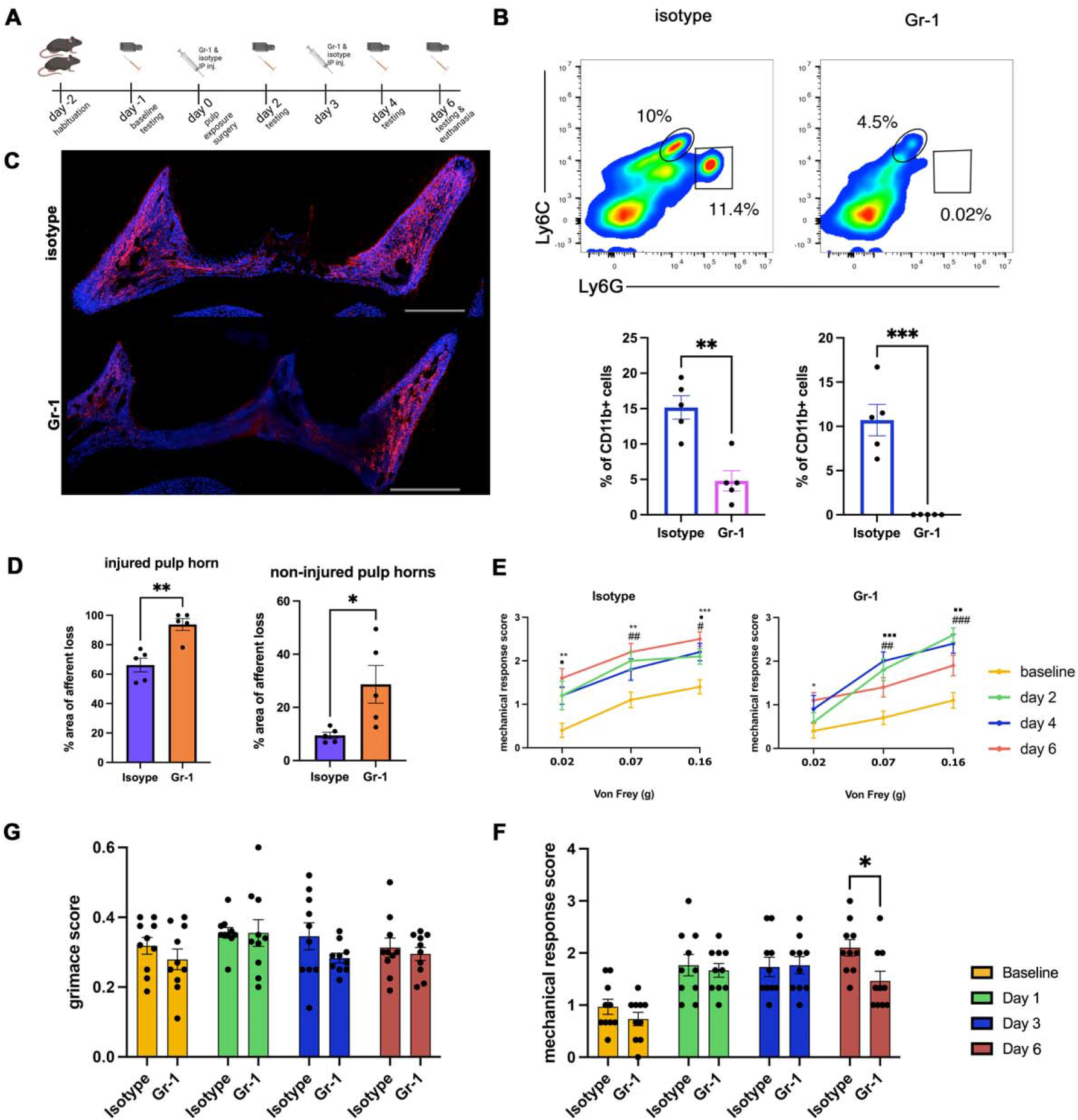
Neutrophil/monocyte depletion (Gr-1) increases sensory afferent loss while leading to delayed reduction in mechanical pain response. A-Schema shows experiment design and timeline for Gr-1 and isotype treatments. B-Flow cytometry representative plots and analysis to validate neutrophil and monocyte depletion using splenocytes. Graphs show % changes of Ly6C high (monocytes) and Ly6G high (neutrophil) cells among total CD11b+ myeloid cells. Unpaired t-test. ∗p < 0.05 ∗∗p < 0.01. Mean ± SEM. C-Representative immunofluorescence staining images showing extent of sensory afferent loss. Red-tuj1, blue-dapi. Scale bar = 200 μm. Original magnification 20X. D-Analysis of extent of sensory afferent loss six days after injury of Gr-1 and isotype treatment groups. In each plot, each point represents an analysis of a section from one animal Unpaired t-tests. ∗p < 0.05 ∗∗p < 0.01. E-The first two plots present mechanical response score of animals of isotype group (n=10) and Gr-1 group (n=10) to different Von Frey filaments at different time points. F-The plot summarizes mechanical response score in isotype and Gr-1 treatment groups at different time points. In this plot, each point represents the mean mechanical response score of one animal obtained by averaging the mechanical response scores given to each Von Frey filament. F-Comparison of grimace scores between animals in isotype vs Gr-1 treatment group at different time points. E-First two plots; repeated measures of ANOVA with Dunnett’s multiple comparisons. The third plot; two-way ANOVA with Sidak’s multiple comparisons test. E-# comparison of day 2 to baseline,▪ □ comparison of day 4 to baseline, * comparison of baseline to day 6. F-Two-way ANOVA with Sidak’s multiple comparison tests. #▪ □ ∗p < 0.05, ##▪ □▪ □ ∗∗p < 0.01, ###▪ □▪ □▪ □ ∗∗∗∗p < 0.0001. Mean ± SEM.

**Figure 6.**
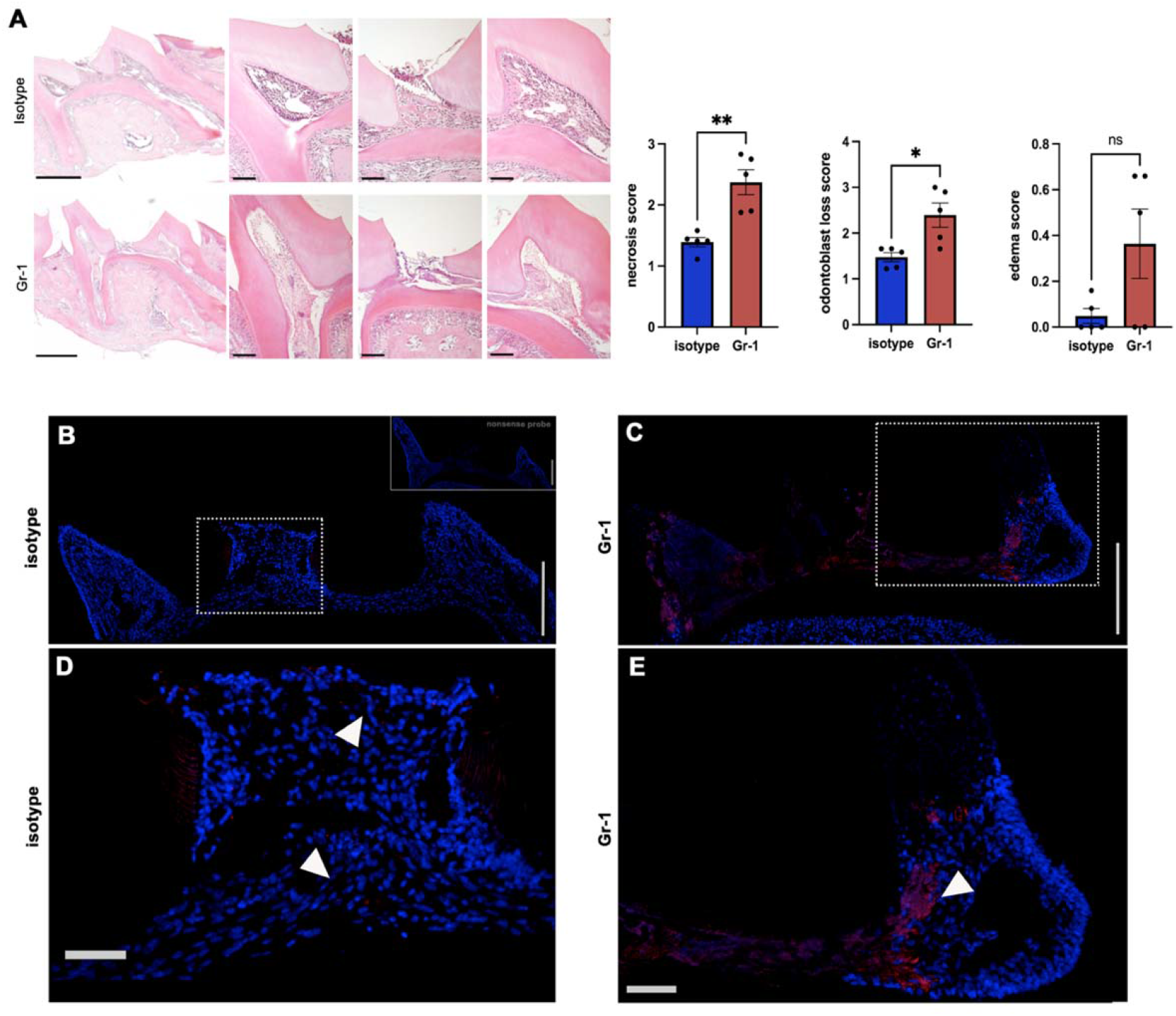
Depletion of neutrophils/monocytes leads to increased tissue necrosis and further bacterial progression. A-Representative H&E staining images and analysis of extent of tissue damage six days after injury of Gr-1 and isotype treatment groups. Scale bar = 500 μm and 100 μm. Original magnification 20X. B-Each point represents an average score obtained from analysis of three sections for one animal. independent/Student’ t-tests. ∗p < 0.05 ∗∗p < 0.01. B-Representative image of 16S FISH from isotype group. The box shows the injured middle pulp with few bacteria present. The image on the top right corner is a section where the nonsense probe was used as a negative control. C-Representative image of 16S FISH from Gr-1 treatment group. The box shows that most of the bacteria are densely present at the injury site but also extends to the non-injured pulp horns. D-Close-up view of middle pulp horn in B, marked by the dotted box. E-Close-up with of the marked area with dotted box in C. Sections are 6 μm. Scale bar = 200 μm and 50 μm. Red-16S, blue-dapi. Original magnification 20X.

**Figure 7.**
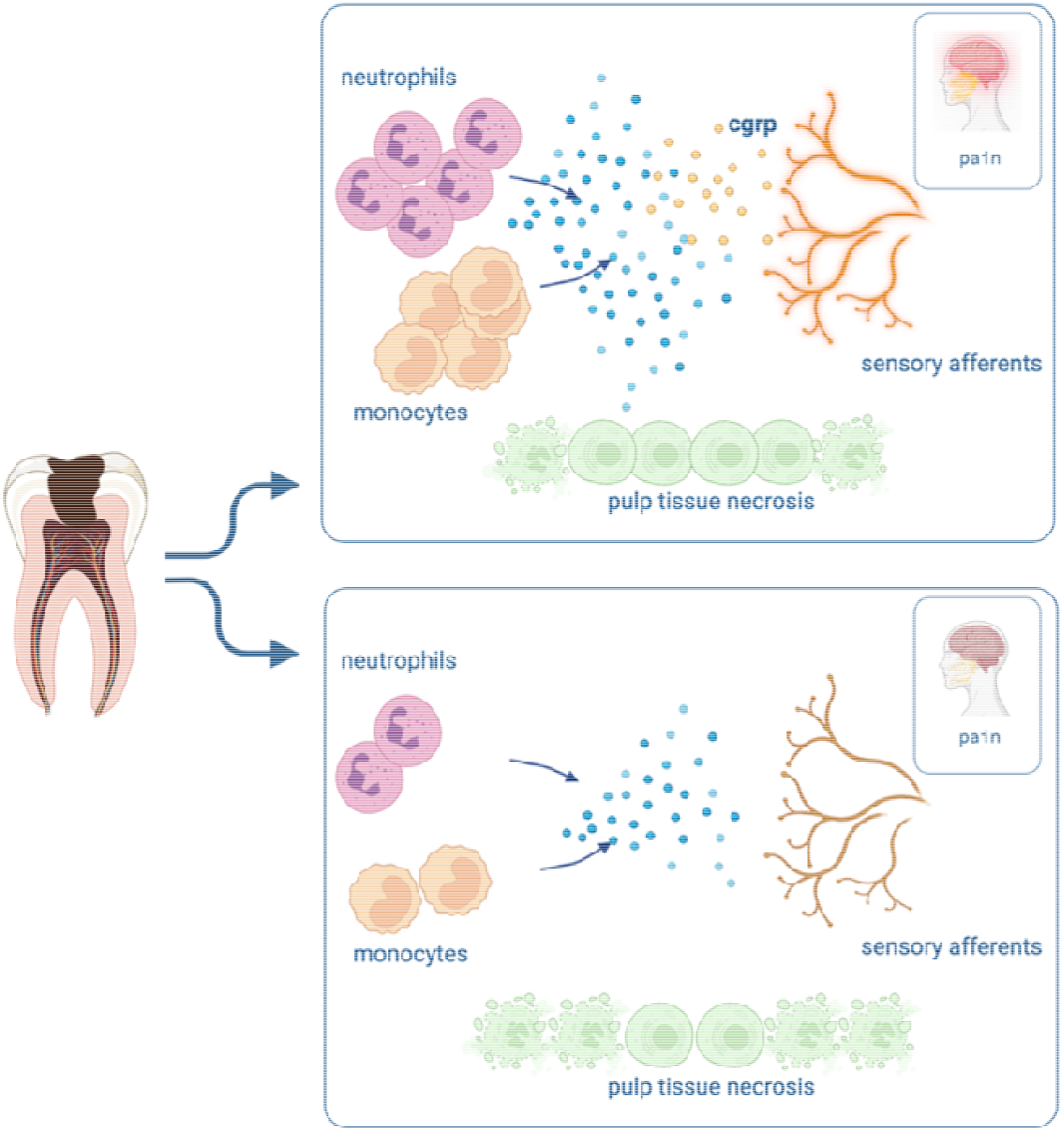
The impact of neuro-immune interactions on pain and tissue necrosis during pulpitis. CGRP modulates innate immune response either by causing vasodilation or by direct involvement in neutrophil recruitment while contributing to spontaneous pain behavior during pulpitis. Neutrophil and monocytes are crucial to protect dental pulp tissue from further bacterial penetration and tissu damage while contributing to mechanical pain response. Prepared using biorender.com

### Depletion of neutrophils and monocytes reduces mechanical hypersensitivity

Finally, we wanted to examine whether reduction in numbers of neutrophils and monocytes altered the pain-like behaviors after dental pulp injury. We saw that the mechanical response score remained similarly high on day 2, 4 and 6 in the isotype group whereas in the Gr-1 group, there was a reduction of mechanical response on day 6 compared to day 2 and 4 (Figure 5-E, F). When we compared the mean mechanical response scores, we found that on day 6, there was a significant difference in mean mechanical response score between isotype and Gr-1 groups. The mean mechanical response score in Gr-1 group was less compared to isotype group (p=0.03) (Figure 5-E, F). We also investigated spontaneous pain-like behavior by grimace scoring, and there was no difference between the two groups across all time points (Figure 5-G). These findings indicate that, when neutrophils and monocytes were significantly reduced, there was more tissue damage, including more sensory afferent loss while there was a reduction in mechanical response scores compared to control group later point (day 6).

## DISCUSSION

Pulpitis can be a debilitating painful experience that is driven by both peripheral and central mechanisms within the trigeminal system [20; 59; 60]. In mice, CGRP injection causes spontaneous pain-like behaviors and activation of CGRP+ fibers increase mechanical sensitivity in mice [22; 30; 43; 56]. By both inducing neuroinflammation at the periphery and sensitization within the trigeminal ganglion, CGRP is likely to play a role in painful pulpitis [2; 34; 47]. In this study we assessed spontaneous pain-like behavior and mechanical sensitivity in calca^−/−^ and calca^+/+^ animals by using grimace scale and facial Von Frey stimulation respectively. We observed at the earliest time point, CGRP contributed to spontaneous pain-like behavior and there was a similar trend at a later time point. Histologically, we saw significant loss of CGRP+ fibers three days after injury which was even more profound seven days after injury. This could be the reason why in this model, the effect of CGRP might have been more evident at the early time point. When we performed facial Von Frey testing we found that there was no difference in mechanical response scores in calca^+/+^ and calca^−/−^ injured animals at different time points. Further studies are needed to investigate mechanisms by which CGRP might be contributing to spontaneous pain-like behavior while not leading to mechanical sensitivity during pulpitis.

CGRP contributed to immune cell recruitment, mostly neutrophils. By investigating the spatial relationship between CGRP+ fibers and neutrophils/monocytes at different time points after injury, we saw that as the innate immune cells were becoming more widespread within the pulp, there was more abundant loss of CGRP+ fibers, especially in the regions densely populated by neutrophils/monocytes. Neutrophils critically perform acute inflammatory duties including bacterial clearance and removal of dying cells, which is protective of tissues. Conversely, neutrophils can contribute to tissue damage in inflammatory diseases such as rheumatoid arthritis (RA) and systemic lupus erythematosus (SLE) [5; 40; 49]. In order to understand whether neutrophils/monocytes are protective or contribute to the progression of tissue damage in pulpitis, we performed a targeted depletion experiment of GR1+ cells. We found that in the absence of continuous recruitment of neutrophils/monocytes, there was increased sensory afferent loss, tissue damage and more bacterial invasion. This suggests that in this model, neutrophils and monocytes are crucial for the clearance of bacteria and protect the pulpal tissue, including the sensory innervation [24; 36; 50].

We found that on day six, there was a rescue of mechanical response scores in neutrophil and monocyte depleted animals [51]. Putting these findings together with our histological examination, we found that in neutrophil/monocyte depleted animals, despite observing more profound sensory afferent die-back and tissue damage, the mechanical nociceptive response was reduced. It is possible that the relationship of pain and sensory afferent loss is complex in pulpitis. In patients experiencing painful pulpitis, there is no clear evidence of a correlation between the severity of pain and the extent of tissue damage [13; 32; 52]. Surprisingly, about 30% of the patients report very mild or even no pain, even though the extent of the pathology is similar to the cases with severe pain [19; 33]. Similarly, in patients with diabetic neuropathy with pathology evidenced by sensory afferent die-back, there may or may not be pain coincident with the sensory disturbances [17; 53]. The mechanisms that could explain how sensory afferent dieback is painful for some patients while it is non-painful for others are not clearly understood in both diseases. It is possible that in our mouse pulpitis model, the sensitization of the nerve fibers driven by cytokines released from neutrophils/monocytes could be a contributor to mechanical sensitivity, whereas die-back of sensory afferents may not contribute to mechanical sensitivity [18; 33]. On the other hand, increased sensory afferent loss when neutrophils were depleted might have led to reduction in mechanical sensory response. Further studies are needed to understand the inflammatory nature of pulpitis pain and the pain due to die-back of sensory afferents, which in some cases, might have a neuropathic component of the pain [7; 10].

## CONCLUSION

In a model of pathogen mediated pulpitis, CGRP release contributed to spontaneous pain-like behavior and neutrophil/monocyte recruitment early in the disease but did not change mechanical hypersensitivity or the progression of pathogen mediated tissue destruction. When neutrophils/monocytes were depleted, there was more sensory afferent die-back and mechanical hypersensitivity responses were lower at the later stage of pulpitis.

## Figures & Legends

**Supplementary Figure 1.**
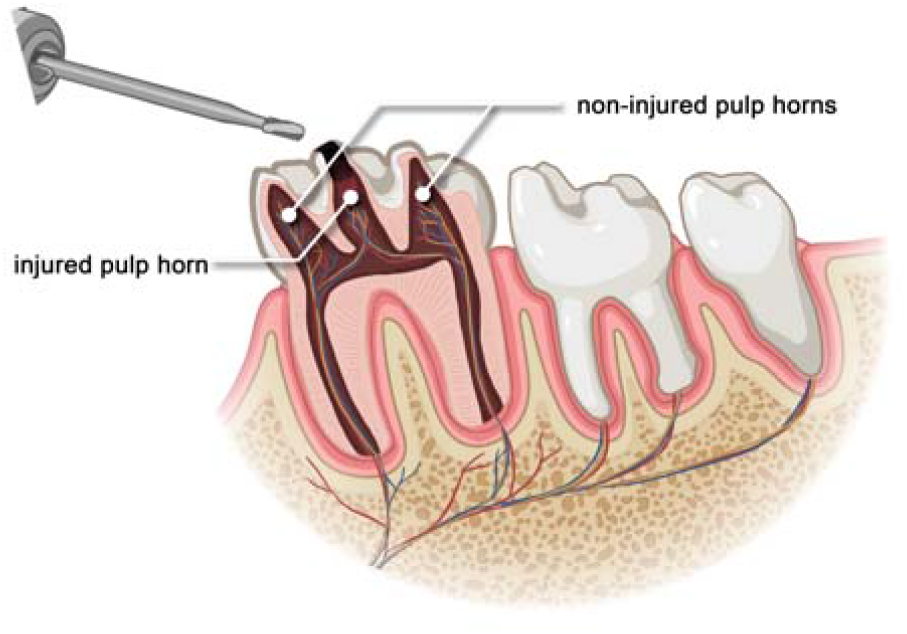
Dental Pulp Injury Model. Representative image of the maxillary first molar tooth. Injury of the middle pulp horn, exposing the pulp to the oral environment is represented. The other two pulp horns are untouched. Prepared by using biorender.com and Procreate.

**Supplementary Figure 2.**
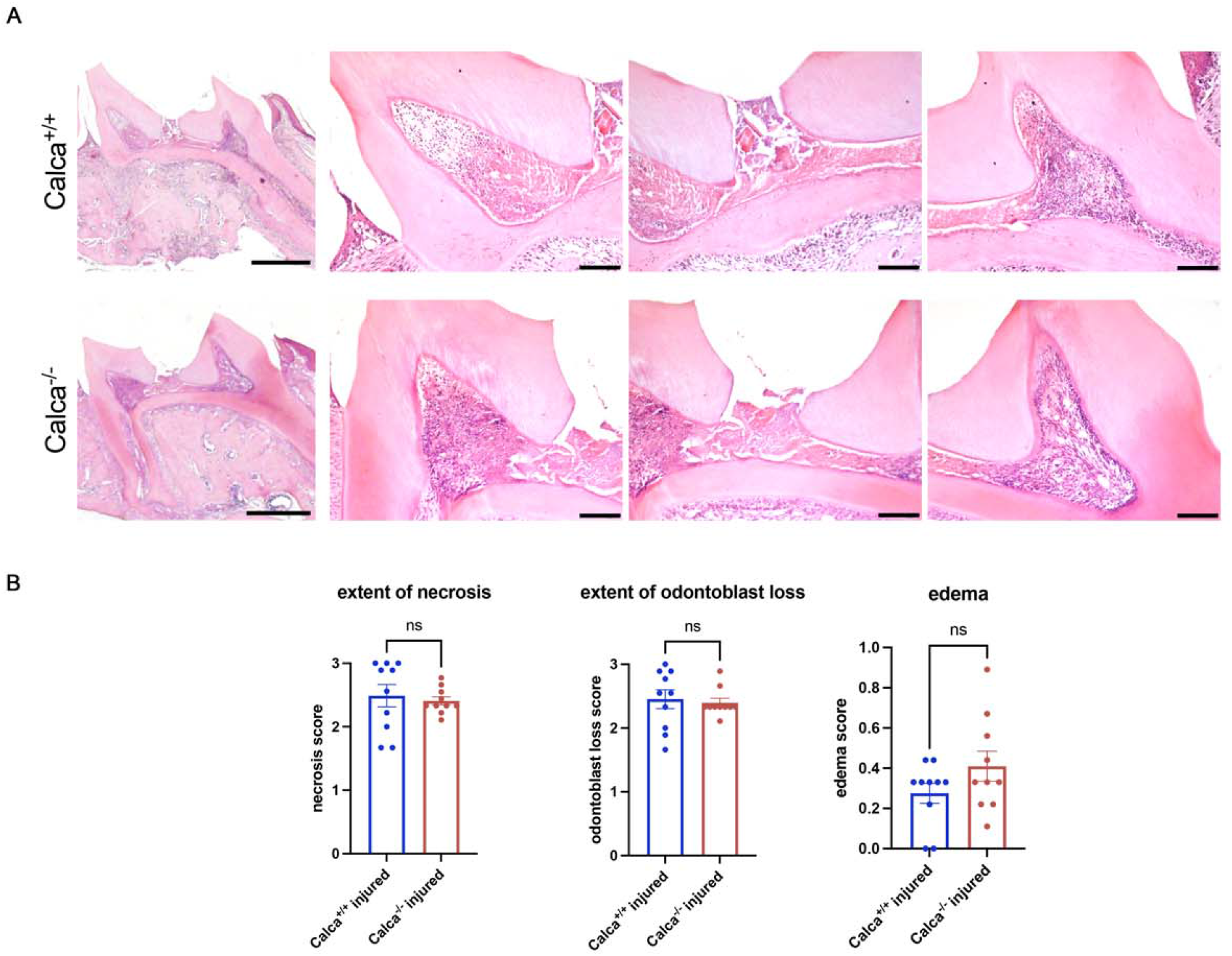
CGRP does not change the tissue damage outcome at seven days after injury. A-Representative H&E staining images and analysis of extent of tissue damage seven days after injury of calca^−/−^ and calca^+/+^ animals. Scale bar = 500 μm and 100 μm. Original magnification 20X. In each graph, each point represents an average score obtained from analysis of three sections for one animal. Unpaired t-tests. ∗p < 0.05.

## REFERENCES

[1] Awawdeh L, Lundy FT, Shaw C, Lamey PJ, Linden GJ, Kennedy JG. Quantitative analysis of substance P, neurokinin A and calcitonin gene-related peptide in pulp tissue from painful and healthy human teeth. Int Endod J 2002;35(1):30–36.

[2] Bletsa A, Fristad I, Berggreen E. Sensory pulpal nerve fibres and trigeminal ganglion neurons express IL-1RI: a potential mechanism for development of inflammatory hyperalgesia. Int Endod J 2009;42(11):978–986.

[3] Byers MR, Taylor PE. Effect of sensory denervation on the response of rat molar pulp to exposure injury. J Dent Res 1993;72(3):613–618.

[4] Carreira EU, Carregaro V, Teixeira MM, Moriconi A, Aramini A, Verri WA, Jr., Ferreira SH, Cunha FQ, Cunha TM. Neutrophils recruited by CXCR1/2 signalling mediate post-incisional pain. Eur J Pain 2013;17(5):654–663.

[5] Chamardani TM, Amiritavassoli S. Inhibition of NETosis for treatment purposes: friend or foe? Mol Cell Biochem 2022;477(3):673–688.

[6] Chu C, Artis D, Chiu IM. Neuro-immune Interactions in the Tissues. Immunity 2020;52(3):464–474.

[7] Costa YM, de Souza PRJ, Marques VAS, Conti PCR, Vivan RR, Duarte MAH, Bonjardim LR. Intraoral Somatosensory Alterations Impact Pulp Sensibility Testing in Patients with Symptomatic Irreversible Pulpitis. J Endod 2020;46(6):786–793.

[8] Cowie AM, Moehring F, O’Hara C, Stucky CL. Optogenetic Inhibition of CGRPalpha Sensory Neurons Reveals Their Distinct Roles in Neuropathic and Incisional Pain. J Neurosci 2018;38(25):5807–5825.

[9] Emrick JJ, von Buchholtz LJ, Ryba NJP. Transcriptomic Classification of Neurons Innervating Teeth. J Dent Res 2020:22034520941837.

[10] Erdogan O, Malek M, Janal MN, Gibbs JL. Sensory testing associates with pain quality descriptors during acute dental pain. Eur J Pain 2019;23(9):1701–1711.

[11] European Society of Endodontology developed b, Duncan HF, Galler KM, Tomson PL, Simon S, El-Karim I, Kundzina R, Krastl G, Dammaschke T, Fransson H, Markvart M, Zehnder M, Bjorndal L. European Society of Endodontology position statement: Management of deep caries and the exposed pulp. Int Endod J 2019;52(7):923–934.

[12] Fristad I, Heyeraas KJ, Kvinnsland IH, Jonsson R. Recruitment of immunocompetent cells after dentinal injuries in innervated and denervated young rat molars: an immunohistochemical study. J Histochem Cytochem 1995;43(9):871–879.

[13] Garfunkel A, Sela J, Ulmansky M. Dental pulp pathosis. Clinicopathologic correlations based on 109 cases. Oral Surg Oral Med Oral Pathol 1973;35(1):110–117.

[14] Ghasemlou N, Chiu IM, Julien JP, Woolf CJ. CD11b+Ly6G-myeloid cells mediate mechanical inflammatory pain hypersensitivity. Proc Natl Acad Sci U S A 2015;112(49):E6808–6817.

[15] Gibbs JL, Melnyk JL, Basbaum AI. Differential TRPV1 and TRPV2 channel expression in dental pulp. J Dent Res 2011;90(6):765–770.

[16] Gibbs JL, Urban R, Basbaum AI. Paradoxical surrogate markers of dental injury-induced pain in the mouse. Pain 2013;154(8):1358–1367.

[17] Gylfadottir SS, Weeracharoenkul D, Andersen ST, Niruthisard S, Suwanwalaikorn S, Jensen TS. Painful and non-painful diabetic polyneuropathy: Clinical characteristics and diagnostic issues. J Diabetes Investig 2019;10(5):1148–1157.

[18] Hall BE, Zhang L, Sun ZJ, Utreras E, Prochazkova M, Cho A, Terse A, Arany P, Dolan JC, Schmidt BL, Kulkarni AB. Conditional TNF-alpha Overexpression in the Tooth and Alveolar Bone Results in Painful Pulpitis and Osteitis. J Dent Res 2016;95(2):188–195.

[19] Hasler JE, Mitchell DF. Painless pulpitis. J Am Dent Assoc 1970;81(3):671–677.

[20] Henry MA, Hargreaves KM. Peripheral mechanisms of odontogenic pain. Dent Clin North Am 2007;51(1):19-44, v.

[21] Hohlbaum K, Corte GM, Humpenoder M, Merle R, Thone-Reineke C. Reliability of the Mouse Grimace Scale in C57BL/6JRj Mice. Animals (Basel) 2020;10(9).

[22] Iyengar S, Johnson KW, Ossipov MH, Aurora SK. CGRP and the Trigeminal System in Migraine. Headache 2019;59(5):659–681.

[23] Iyengar S, Ossipov MH, Johnson KW. The role of calcitonin gene-related peptide in peripheral and central pain mechanisms including migraine. Pain 2017;158(4):543–559.

[24] Kim MH, Liu W, Borjesson DL, Curry FR, Miller LS, Cheung AL, Liu FT, Isseroff RR, Simon SI. Dynamics of neutrophil infiltration during cutaneous wound healing and infection using fluorescence imaging. J Invest Dermatol 2008;128(7):1812–1820.

[25] Kolaczkowska E, Kubes P. Neutrophil recruitment and function in health and inflammation. Nat Rev Immunol 2013;13(3):159–175.

[26] Langford DJ, Bailey AL, Chanda ML, Clarke SE, Drummond TE, Echols S, Glick S, Ingrao J, Klassen-Ross T, Lacroix-Fralish ML, Matsumiya L, Sorge RE, Sotocinal SG, Tabaka JM, Wong D, van den Maagdenberg AM, Ferrari MD, Craig KD, Mogil JS. Coding of facial expressions of pain in the laboratory mouse. Nat Methods 2010;7(6):447–449.

[27] Lee C, Ramsey A, De Brito-Gariepy H, Michot B, Podborits E, Melnyk J, Gibbs JLG. Molecular, cellular and behavioral changes associated with pathological pain signaling occur after dental pulp injury. Molecular pain 2017;13:1744806917715173.

[28] Levin LG, Law AS, Holland GR, Abbott PV, Roda RS. Identify and define all diagnostic terms for pulpal health and disease states. J Endod 2009;35(12):1645–1657.

[29] Löken L, Etlin A, Bernstein M, Steyert M, Kuhn J, Hamel K, Llewellyn-Smith I, Braz J, Basbaum A. Dorsal horn CGRP-expressing interneurons contribute to nerve injury-induced mechanical hypersensitivity. bioRxiv 2020:2020.2006.2008.131839.

[30] Marquez de Prado B, Hammond DL, Russo AF. Genetic enhancement of calcitonin gene-related Peptide-induced central sensitization to mechanical stimuli in mice. J Pain 2009;10(9):992–1000.

[31] McCoy ES, Taylor-Blake B, Street SE, Pribisko AL, Zheng J, Zylka MJ. Peptidergic CGRPalpha primary sensory neurons encode heat and itch and tonically suppress sensitivity to cold. Neuron 2013;78(1):138–151.

[32] Mejare IA, Axelsson S, Davidson T, Frisk F, Hakeberg M, Kvist T, Norlund A, Petersson A, Portenier I, Sandberg H, Tranaeus S, Bergenholtz G. Diagnosis of the condition of the dental pulp: a systematic review. Int Endod J 2012;45(7):597–613.

[33] Michaelson PL, Holland GR. Is pulpitis painful? Int Endod J 2002;35(10):829–832.

[34] Michot B, Casey SM, Gibbs JL. Effects of CGRP-Primed Dental Pulp Stem Cells on Trigeminal Sensory Neurons. J Dent Res 2021;100(11):1273–1280.

[35] Nadeau S, Filali M, Zhang J, Kerr BJ, Rivest S, Soulet D, Iwakura Y, de Rivero Vaccari JP, Keane RW, Lacroix S. Functional recovery after peripheral nerve injury is dependent on the pro-inflammatory cytokines IL-1beta and TNF: implications for neuropathic pain. J Neurosci 2011;31(35):12533–12542.

[36] Nakamura K, Yamasaki M, Nishigaki N, Iwama A, Imaizumi I, Nakamura H, Kameyama Y. Effect of methotrexate-induced neutropenia on pulpal inflammation in rats. J Endod 2002;28(4):287–290.

[37] Oh-hashi Y, Shindo T, Kurihara Y, Imai T, Wang Y, Morita H, Imai Y, Kayaba Y, Nishimatsu H, Suematsu Y, Hirata Y, Yazaki Y, Nagai R, Kuwaki T, Kurihara H. Elevated sympathetic nervous activity in mice deficient in alphaCGRP. Circ Res 2001;89(11):983–990.

[38] Parisien M, Lima LV, Dagostino C, El-Hachem N, Drury GL, Grant AV, Huising J, Verma V, Meloto CB, Silva JR, Dutra GGS, Markova T, Dang H, Tessier PA, Slade GD, Nackley AG, Ghasemlou N, Mogil JS, Allegri M, Diatchenko L. Acute inflammatory response via neutrophil activation protects against the development of chronic pain. Sci Transl Med 2022;14(644):eabj9954.

[39] Park SK, Choi SK, Kim YG, Choi SY, Kim JW, Seo SH, Lee JH, Bae YC. Parvalbumin-, substance P- and calcitonin gene-related peptide-immunopositive axons in the human dental pulp differ in their distribution of varicosities. Sci Rep 2020;10(1):10672.

[40] Perez-Figueroa E, Alvarez-Carrasco P, Ortega E, Maldonado-Bernal C. Neutrophils: Many Ways to Die. Front Immunol 2021;12:631821.

[41] Pinho-Ribeiro FA, Baddal B, Haarsma R, O’Seaghdha M, Yang NJ, Blake KJ, Portley M, Verri WA, Dale JB, Wessels MR, Chiu IM. Blocking Neuronal Signaling to Immune Cells Treats Streptococcal Invasive Infection. Cell 2018;173(5):1083–1097 e1022.

[42] Pinho-Ribeiro FA, Verri WA, Jr., Chiu IM. Nociceptor Sensory Neuron-Immune Interactions in Pain and Inflammation. Trends Immunol 2017;38(1):5–19.

[43] Rea BJ, Wattiez AS, Waite JS, Castonguay WC, Schmidt CM, Fairbanks AM, Robertson BR, Brown CJ, Mason BN, Moldovan-Loomis MC, Garcia-Martinez LF, Poolman P, Ledolter J, Kardon RH, Sowers LP, Russo AF. Peripherally administered calcitonin gene-related peptide induces spontaneous pain in mice: implications for migraine. Pain 2018;159(11):2306–2317.

[44] Ren K, Dubner R. Interactions between the immune and nervous systems in pain. Nat Med 2010;16(11):1267–1276.

[45] Rossi HL, See LP, Foster W, Pitake S, Gibbs J, Schmidt B, Mitchell CH, Abdus-Saboor I. Evoked and spontaneous pain assessment during tooth pulp injury. Sci Rep 2020;10(1):2759.

[46] Russell FA, King R, Smillie SJ, Kodji X, Brain SD. Calcitonin gene-related peptide: physiology and pathophysiology. Physiol Rev 2014;94(4):1099–1142.

[47] Sattari M, Mozayeni MA, Matloob A, Mozayeni M, Javaheri HH. Substance P and CGRP expression in dental pulps with irreversible pulpitis. Aust Endod J 2010;36(2):59–63.

[48] Schindelin J, Arganda-Carreras I, Frise E, Kaynig V, Longair M, Pietzsch T, Preibisch S, Rueden C, Saalfeld S, Schmid B, Tinevez JY, White DJ, Hartenstein V, Eliceiri K, Tomancak P, Cardona A. Fiji: an open-source platform for biological-image analysis. Nat Methods 2012;9(7):676–682.

[49] Schultz BM, Acevedo OA, Kalergis AM, Bueno SM. Role of Extracellular Trap Release During Bacterial and Viral Infection. Front Microbiol 2022;13:798853.

[50] Segal AW. How neutrophils kill microbes. Annu Rev Immunol 2005;23:197–223.

[51] Segelcke D, Pradier B, Reichl S, Schafer LC, Pogatzki-Zahn EM. Investigating the Role of Ly6G(+) Neutrophils in Incisional and Inflammatory Pain by Multidimensional Pain-Related Behavioral Assessments: Bridging the Translational Gap. Front Pain Res (Lausanne) 2021;2:735838.

[52] Seltzer S, Bender IB, Ziontz M. The dynamics of pulp inflammation: correlations between diagnostic data and actual histologic findings in the pulp. Oral Surg Oral Med Oral Pathol 1963;16:846–871 contd.

[53] Spallone V, Greco C. Painful and painless diabetic neuropathy: one disease or two? Curr Diab Rep 2013;13(4):533–549.

[54] Usoskin D, Furlan A, Islam S, Abdo H, Lonnerberg P, Lou D, Hjerling-Leffler J, Haeggstrom J, Kharchenko O, Kharchenko PV, Linnarsson S, Ernfors P. Unbiased classification of sensory neuron types by large-scale single-cell RNA sequencing. Nat Neurosci 2015;18(1):145–153.

[55] Wang Y, Bao M, Hou C, Wang Y, Zheng L, Peng Y. The Role of TNF-alpha in the Pathogenesis of Temporomandibular Disorders. Biol Pharm Bull 2021;44(12):1801–1809.

[56] Wattiez AS, Gaul OJ, Kuburas A, Zorrilla E, Waite JS, Mason BN, Castonguay WC, Wang M, Robertson BR, Russo AF. CGRP induces migraine-like symptoms in mice during both the active and inactive phases. J Headache Pain 2021;22(1):62.

[57] Wattiez AS, Wang M, Russo AF. CGRP in Animal Models of Migraine. Handb Exp Pharmacol 2019;255:85–107.

[58] Whittaker AL, Liu Y, Barker TH. Methods Used and Application of the Mouse Grimace Scale in Biomedical Research 10 Years on: A Scoping Review. Animals (Basel) 2021;11(3).

[59] Worsley MA, Allen CE, Billinton A, King AE, Boissonade FM. Chronic tooth pulp inflammation induces persistent expression of phosphorylated ERK (pERK) and phosphorylated p38 (pp38) in trigeminal subnucleus caudalis. Neuroscience 2014;269:318–330.

[60] Zhang S, Chiang CY, Xie YF, Park SJ, Lu Y, Hu JW, Dostrovsky JO, Sessle BJ. Central sensitization in thalamic nociceptive neurons induced by mustard oil application to rat molar tooth pulp. Neuroscience 2006;142(3):833–842.

